# Autism- and epilepsy-associated *EEF1A2* mutations lead to translational dysfunction and altered actin bundling

**DOI:** 10.1101/2023.06.06.543912

**Authors:** Muhaned S. Mohamed, Eric Klann

## Abstract

Protein synthesis is a fundamental cellular process in neurons that is essential for synaptic plasticity and memory consolidation. Here, we describe our investigations of a neuron- and muscle-specific translation factor, <u>e</u>ukaryotic <u>E</u>longation <u>F</u>actor <u>1a2</u> (eEF1A2), which when mutated in patients results in autism, epilepsy, and intellectual disability. We characterize three most common *EEF1A2* patient mutations, G70S, E122K, and D252H, and demonstrate that all three mutations decrease *de novo* protein synthesis and elongation rates in HEK293 cells. In mouse cortical neurons, the *EEF1A2* mutations not only decrease *de novo* protein synthesis, but also alter neuronal morphology, regardless of endogenous levels of eEF1A2, indicating that the mutations act via a toxic gain of function. We also show that eEF1A2 mutant proteins display increased tRNA binding and decreased actin bundling activity, suggesting that these mutations disrupt neuronal function by decreasing tRNA availability and altering the actin cytoskeleton. More broadly, our findings are consistent with the idea that eEF1A2 acts as a bridge between translation and the actin skeleton, which is essential for proper neuron development and function.

**Significance Statement:** <u>E</u>ukaryotic <u>E</u>longation <u>F</u>actor 1A2 (eEF1A2) is a muscle- and neuron-specific translation factor responsible for bringing charge tRNAs to the elongating ribosome. Why neurons express this unique translation factor is unclear; however, it is known that mutations in *EEF1A2* cause severe drug-resistant epilepsy, autism and neurodevelopmental delay. Here, we characterize the impact of three common disease-causing mutations in *EEF1A2* and demonstrate that they cause decreased protein synthesis via reduced translation elongation, increased tRNA binding, decreased actin bundling activity, as well as altered neuronal morphology. We posit that eEF1A2 serves as a bridge between translation and the actin cytoskeleton, linking these two processes that are essential for neuronal function and plasticity.

## INTRODUCTION

Autism spectrum disorders (ASDs) are a class of complex neurodevelopmental disorders with a wide variety of neurological comorbidities including epilepsy, intellectual disability, attention deficit hyperactivity disorder (ADHD), and sleep disturbances (1). Although there are many genes associated with ASD, a number of genetic aberrations are found in genes that regulate neuronal protein synthesis, such as *FMR1*, which encodes fragile X messenger ribonuclear protein 1 and causes fragile X syndrome, and *TSC2*, which encodes tuberin and causes tuberous sclerosis complex (2). Neuronal translational control has been most intensely studied at the initiation step by signaling pathways that control the activity of initiation factors such <u>e</u>ukaryotic Initiation <u>F</u>actor 2α(eIF2α and <u>e</u>ukaryotic Initiation <u>F</u>actor <u>4E</u>-<u>b</u>inding <u>p</u>rotein (4E-BP) (3). However, there is a growing body evidence that regulation of protein synthesis at the elongation step is critical for neuronal function and that neurons preferentially translate certain mRNAs by controlling elongation (4). Consistent with this observation, neurons express a unique elongation factor isoform, <u>e</u>ukaryotic <u>E</u>longation <u>F</u>actor <u>1α2</u> (eEF1A2), which replaces the ubiquitously expressed isoform <u>e</u>ukaryotic <u>E</u>longation <u>F</u>actor <u>1α1</u> (eEF1A1) as neurons develop and becomes the predominant factor (5). *De novo* mutations in *EEF1A2* have been found in patients with severe and often drug-resistant epilepsy, autism, and intellectual disability (6-8). Three of the most common disease-causing mutations in *EEF1A2* found in patients with severe autism and epilepsy are G70S, E122K, D252H. It is not known how these disease-associated mutations affect protein synthesis and neuronal function.

eEF1A is responsible for carrying out first step eukaryotic translation elongation by delivering amino-acylated tRNAs to the A-site of the ribosome in a GTP-dependent manner. Following codon-anticodon recognition, GTP is hydrolyzed and eEF1A is released from the ribosome (9). eEF1A requires a guanine nucleotide exchange factor (GEF) to exchange GDP for GTP in order to bind another tRNA. The eEF1A GEF is composed of three subunits, eEF1B2, eEF1D, and eEF1G, which together compose the eEF1 (<u>e</u>ukaryotic <u>E</u>longation <u>F</u>actor 1) complex (5). Without the eEF1 complex, GDP exchange on eEF1A is quite slow; there is a nearly 700-fold increase in exchange rate in the presence of eEF1B2 (10). eEF1A proteins also have been shown to bind and bundle actin, with the A1 and A2 isoforms having different activities despite being nearly 93% identical and 96% similar on the amino acid level (11, 12). In mouse neurons, eEF1A2 is critical for survival; deletion of *Eef1a2* in a mouse model known as the *wasted* mouse results in muscle wasting, gait abnormalities, and death shortly after weaning (13). The emergence of these phenotypes corresponds with a developmental switch in isoform expression where the expression of the eEF1A2 isoform increases and expression of the eEF1A1 isoform decreases. This isoform switch occurs around postnatal day 14 in mice and shortly after birth in humans (14).

It is unclear why neurons require a unique translation elongation factor for a universal process such as protein synthesis. The neurological abnormalities associated with *EEF1A2* mutations suggest it plays an important role in the development and activity of neurons. Other published evidence also suggests the eEF1A2 isoform plays a unique role in neurons. Despite the similarities between the two isoforms, it has been shown that the overexpression of eEF1A2, but not eEF1A1, in neurons increases *de novo* protein synthesis (15). It also has been shown that the eEF1A2 is phosphorylated in response to neuronal stimulation, thereby triggering both its dissociation from the guanine exchange factor (GEF) eEF1B2 and decreased association with actin (16). Given this evidence we set out to characterize the effects of autism- and epilepsy-associated mutations in *EEF1A2* in HEK293 cells and mouse cortical neurons. We found that the disease-associated mutations in *EEF1A2* disrupt protein synthesis by decreasing elongation rates and alter the morphological development of cortical neurons. We also demonstrate that the disease-causing mutations *EEF1A2* increase tRNA binding and decrease actin bundling activity, suggesting that the changes in neuronal protein synthesis and neuronal morphology are due to these respective changes to eEF1A2 function. Our results indicate that eEF1A2 serves a dual role in neurons, both maintaining protein synthesis and contributing to cytoskeletal organization.

## RESULTS

### *EEF1A2* mutations decrease de novo protein synthesis and elongation rates in HEK293 cells

The three most common *EEF1A2* mutations exist in separate functional domains of eEF1A2, where the G70S mutation lies in the GTP binding domain, the E122K mutation lies in the tRNA binding domain and the D252H mutation is near the actin binding domain. Yet recent structural studies suggest that both the G70 and D252 residues may be directly involve in interacting with ribosome and assist in proofreading (Fig. 1A)(17). Given the central role of eEF1A2 in translation, we set out to determine whether disease-associated mutations in *EEF1A2* alter de novo protein synthesis. HEK293 cells, which express very low levels of eEF1A2, were transiently transfected with either wild-type eEF1A2 or the G70S, E122K, D252H mutants for 24 hours followed by a SUnSET assay to measure de novo protein synthesis (18, 19). We found that G70S and E122K mutations exhibited an approximately 40% reduction in *de novo* protein synthesis whereas the D252H mutation showed a more moderate reduction in *de novo* protein synthesis (Fig. 1B-C). The decrease in protein synthesis caused by the G70S, E122K and D252H mutations was not due to activation of the integrated stress response, as we did not observe any alteration in the phosphorylation of eIF2α or the levels of its downstream effector ATF4 (Fig. S1A). This suggests that the decrease in protein synthesis is due to an effect of the mutant protein on translation but not via a cellular stress response.

**Figure 1.**
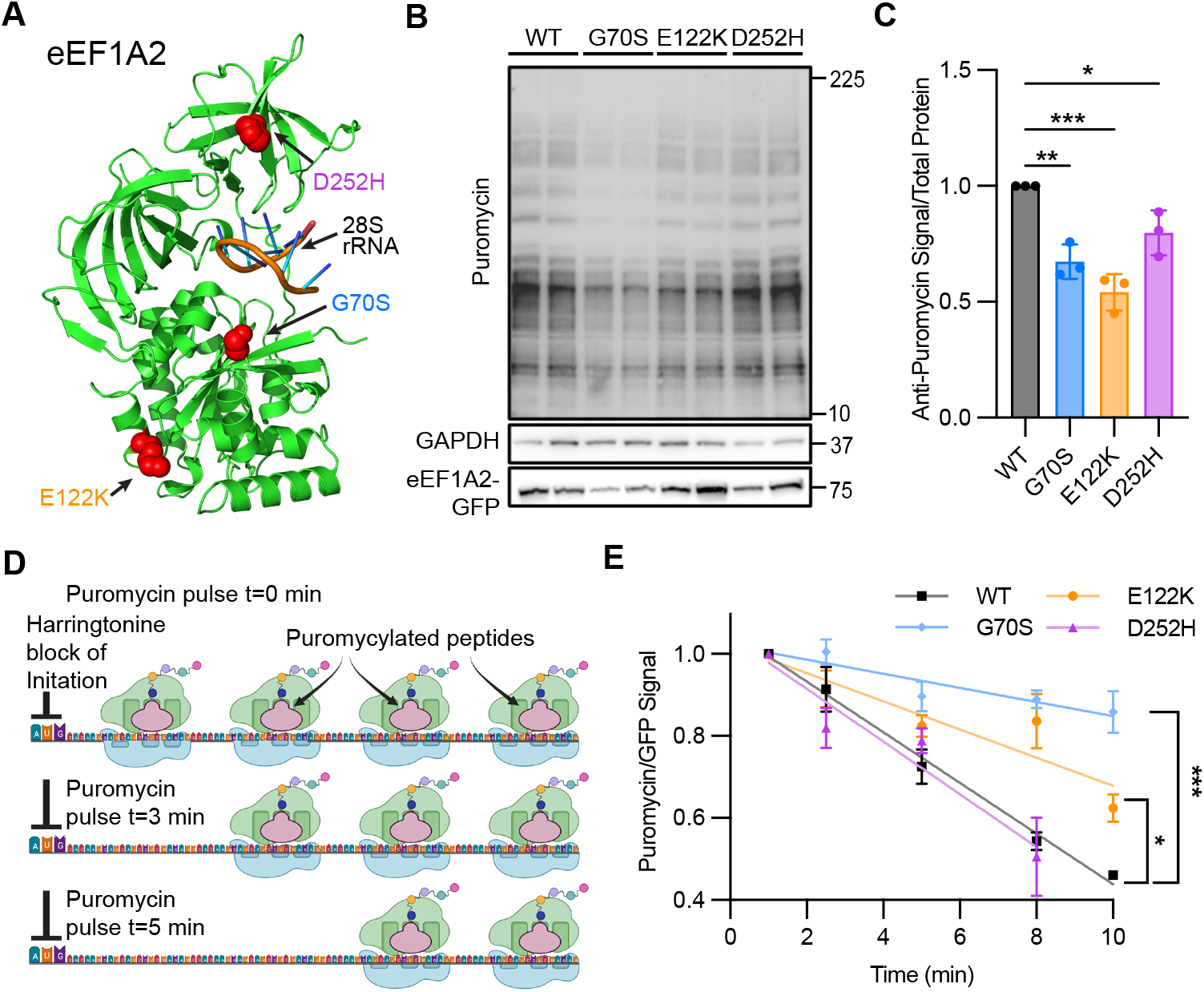
A) *EEF1A2* mutations decrease de novo protein synthesis and elongation rate in HEK293 cells. eEF1A2 Cryo-EM structure (PDB: 8B6Z) interacting with the mammalian ribosome (28s rRNA). Red labeled residues are the autism and-epilepsy causing mutations. B) Representative western blot of SUnSET assay in HEK293 cells transfected with wild-type *EEF1A2*, or G70S, E122K, D252H mutants. C) Quantification of three independent experiments of SUnSET assay in HEK293 cells. Data in (B) are means ± standard deviation (SD) and are normalized to wild type *EEF1A2*. One way ANOVA with Dunnet test **P=0.0015, ***P=0.0002, *P=0.0276 D) Schematic of SuNRISE Assay E) SuNRISE assay in HEK293 cells transfected with wild-type *EEF1A2*, or G70S, E122K, D252H mutants. Data points are means of 4 replicates ± standard error of the mean (SEM). One-way ANOVA with Tukey’s multiple comparison test *P<0.03, **P<0.009, ***P<0.0001

Because eEF1A2 catalyzes the first step of translation elongation, we measured the rate of elongation in transfected HEK293 cells using SUnRISE (20), which quantifies elongation rates by measuring the decrease in puromycin labeled peptides due to ribosome run-off after blocking translation initiation with harringtonine (Fig. 1D). We first performed control experiments and found that blocking ribosome translocation with emetine did not cause a decrease in the puromycin signal, which indicates that there was no ribosome run-off (Fig. S1B). We proceeded to examine the impact of the *EEF1A2* mutations on elongation. Consistent with the results of the SUnSET experiments, we found that the G70S and E122K *EEF1A2* mutations decreased the rate of elongation, whereas the D252H mutation did not decrease the elongation rate (Fig. 1E). In all, the G70S, E122K, and D252H mutations decrease de novo protein synthesis in HEK293 cells most likely be decreasing the rate of elongation.

### *EEF1A2* mutations decrease de novo protein synthesis and alter neuronal morphology in primary mouse cortical neurons

To determine whether *EEF1A2* mutations act via gain of function, loss of function, or a dominant negative effect, we isolated primary cortical neurons from embryonic day 16 pups that were either *Eef1a2* null, *Eef1a2* heterozygous, or wild type. We hypothesized that *EEF1A2* mutations act via a gain of function and would alter protein synthesis in neurons derived from all three types of mice. To measure de novo protein synthesis, neurons were transfected with either wild-type *EEF1A2* or the G70S, E122K, D252H mutants at DIV 6 and labeled with puromycin for SUnSET at DIV14 (Fig. 2A). We found that overexpression of eEF1A2 increased overall protein synthesis in all of the neurons regardless of endogenous eEF1A2 expression (Fig 2B). Conversely, the *EEF1A2* mutants did not increase protein synthesis, but instead reduced it below baseline levels (Fig. 2C-E). This suggests that eEF1A2 mutants act to reduced de novo protein synthesis via a toxic gain of function mechanism.

**Figure 2.**
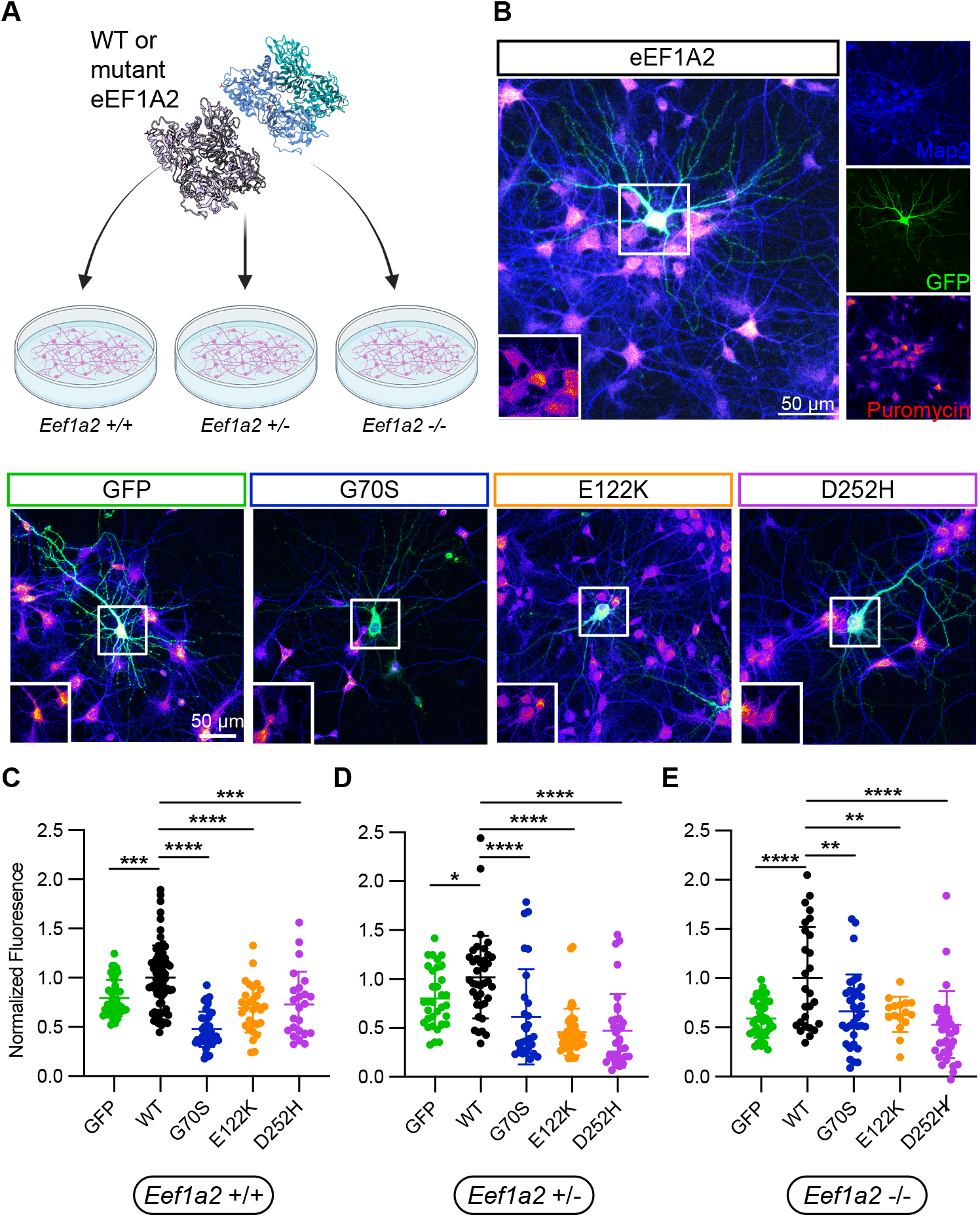
*EEF1A2* mutations decrease de novo protein synthesis in primary mouse cortical neurons A) Schematic of experiment paradigm Representative images of B) SUnSET Assay in *Eef1a2* +/+ neurons transfected with either GFP, wild-type *EEF1A2*, or G70S, E122K, D252H mutants at DIV 14. Insets show puromycin signal from transfected neuron soma. C) SUnSET Assay in *Eef1a2* +/+ neurons transfected with either GFP (n=40 neurons), wild-type *EEF1A2* (n=73), or G70S (n=44), E122K (n=29), D252H (n=24) mutants at DIV 14. Single neuron data (dots) are plotted and mean ± SEM are as bars. Tukey’s multiple comparison test **P<0.002, *** P<0.0002 D) SUnSET Assay in *Eef1a*2 +/-neurons transfected with either GFP (n=33), wild-type *EEF1A2* (n=38), or G70S (n=29), E122K (n=42), D252H (n=35) mutants at DIV 14. Single neuron data (dots) are plotted and mean ± SEM are as bars. Tukey’s multiple comparison test *P<0.05, *** P<0.0001 E) SUnSET Assay in *Eef1a2* -/- neurons transfected with either GFP (n=42), wild-type EEF1A2 (n=29), or G70S (n=41), E122K (n=34), D252H (n=33) mutants at DIV 14. Single neuron data (dots) are plotted and mean ± SEM are as bars. Tukey’s multiple comparison test **P<0.009, **** P<0.0001

Neurons in culture exhibit a similar developmental switch in eEF1A isoforms as was observed in the developing brain (14, 16), which was reproduced in our neuronal cultures (Fig. S2A-B). Interestingly this switch in eEF1A expression also corresponds to an increase in de novo protein synthesis as neurons develop (Fig. S2C). At baseline, complete loss of *Eef1a2* expression in mice is lethal after weaning. In contrast, mice with loss of only one *Eef1a2* allele develop normally and exhibit no physical or neurological deficits (21). Given the severe neurodevelopmental delay observed in patients with *EEF1A2* mutations, we postulated that the mutations would affect dendritic development and arborization. To explore this possibility, we transfected *Eef1a2* null, *Eef1a2* heterozygous, or wild-type primary cortical neurons with mCherry and either *EEF1A2* or the G70S, E122K, D252H mutants at DIV 6 and fixed the neurons at DIV14 for Sholl analysis (Fig 3A). We found no difference in dendritic arborization and morphology between wild-type and *Eef1a2* heterozygous neurons, but a small decrease in arborization of *Eef1a2* null neurons (Fig. S3). Similarly, overexpression of eEF1A2 did not significantly alter dendritic arborization. However, the expression of *EEF1A2* mutants greatly reduced the number, branching, and length of dendrites in all neurons regardless of the levels of endogenous eEF1A2 expression (Figure 3B-D). However, we did note that in the wild type and *Eef1a2* null neurons the E122K and D252H mutants respectively did not reduce dendritic arborization significantly. This is most likely due to slight differences in the effects of individual mutations and variations in expression of transfected plasmids. Overall, these results suggest that the disease-causing *EEF1A2* mutations act via a toxic gain of function by altering dendritic arborization and outgrowth.

**Figure 3.**
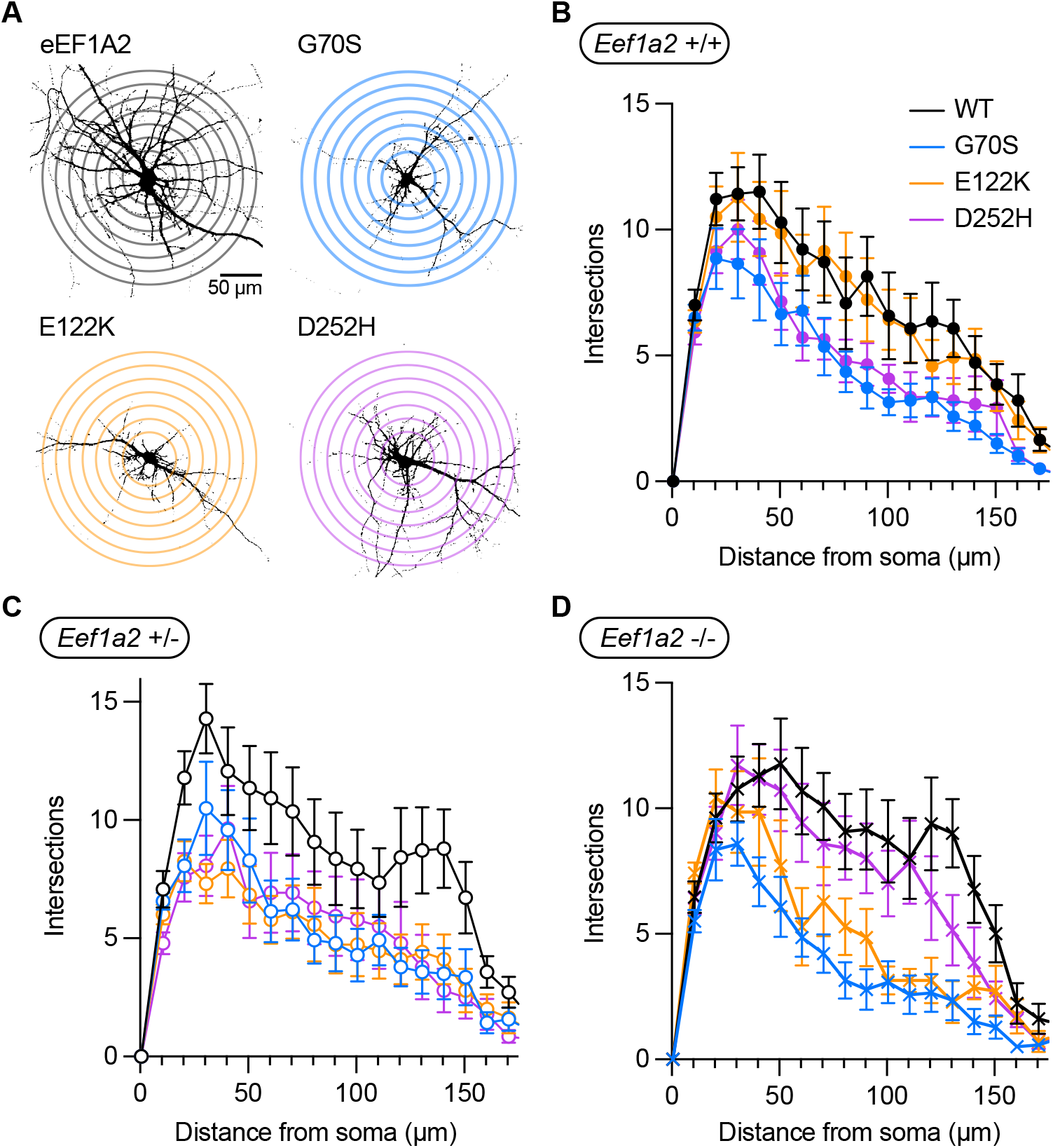
*EEF1A2* mutations alter neuronal morphology in primary mouse cortical neurons A) Representative images of Sholl analysis of *Eef1a2* (+/+) neurons transfected with either wild-type *EEF1A2*, or G70S, E122K, D252H mutants at DIV 14. B) Sholl analysis of *Eef1a2* +/+ neurons comparing (two-way ANOVA) neurons transfected with wild-type *EEF1A2*, with G70S (p<0.0001), E122K (p=0.0712), and D252H (p<0.0001) mutants at DIV 14. Two-way ANOVA with Tukey’s multiple comparison test. Each line graph represents the number of dendritic intersections with concentric shells drawn at 10um from the center of the soma at 10um intervals. C) Sholl analysis of *Eef1a2* +/-neurons comparing neurons transfected with wild-type *EEF1A2*, with G70S (p<0.0001), E122K (p<0.0001), and D252H (p<0.0001) mutants at DIV 14. Two-way ANOVA with Tukey’s multiple comparison test D) Sholl analysis of *Eef1a2* -/-neurons comparing neurons transfected with wild-type *EEF1A2*, with G70S (p<0.0001), E122K (p<0.0001), and D252H (p=0.068) mutants at DIV 14. Two-way ANOVA with Tukey’s multiple comparison test.

### *EEF1A2* mutations alter tRNA binding and actin bundling activity

To carry out its canonical function as elongation factor, eEF1A2 needs to bind tRNA, followed by hydrolysis of GTP, and then interact with the eEF1 complex to exchange GDP to GTP (9). Noncanonically, eEF1A2 also binds and bundle actin as a homodimer (Figure 4A) (22). Therefore, we determined which of the known functions of eEF1A2 were perturbed by the disease-causing mutations. Despite the fact that the G70S and E122K mutations occur near the eEF1A2 active site, we found no changes in intrinsic GTPase activity by these or the D252H mutation (Figure S4A). However, the E122K and D252H mutations were found to significantly increase tRNA binding (Figure 4B-C), suggesting that either the eEF1A2 mutant proteins are sequestering tRNAs or they preferentially exist in the monomer tRNA-bound state rather than the dimer state. Given that eEF1A2 requires the eEF1 complex (composed of eEF1B2, eEF1D, eEF1G) to carry out GDP exchange, we preformed co-immunoprecipitations of GFP-tagged eEF1A2 or G70S, E122K, D252H mutants. Only the D252H mutants perturbed eEF1 complex binding (Figure S4B), whereas the G70S and E122K mutants exhibited binding similar to that of WT eEF1A2. These results corroborate previous findings (23) and suggest that the D252H mutant perturbs the EF1 complex because the D252 residue is located in the binding pocket for eEF1B2, (Fig. S4C).

**Figure 4.**
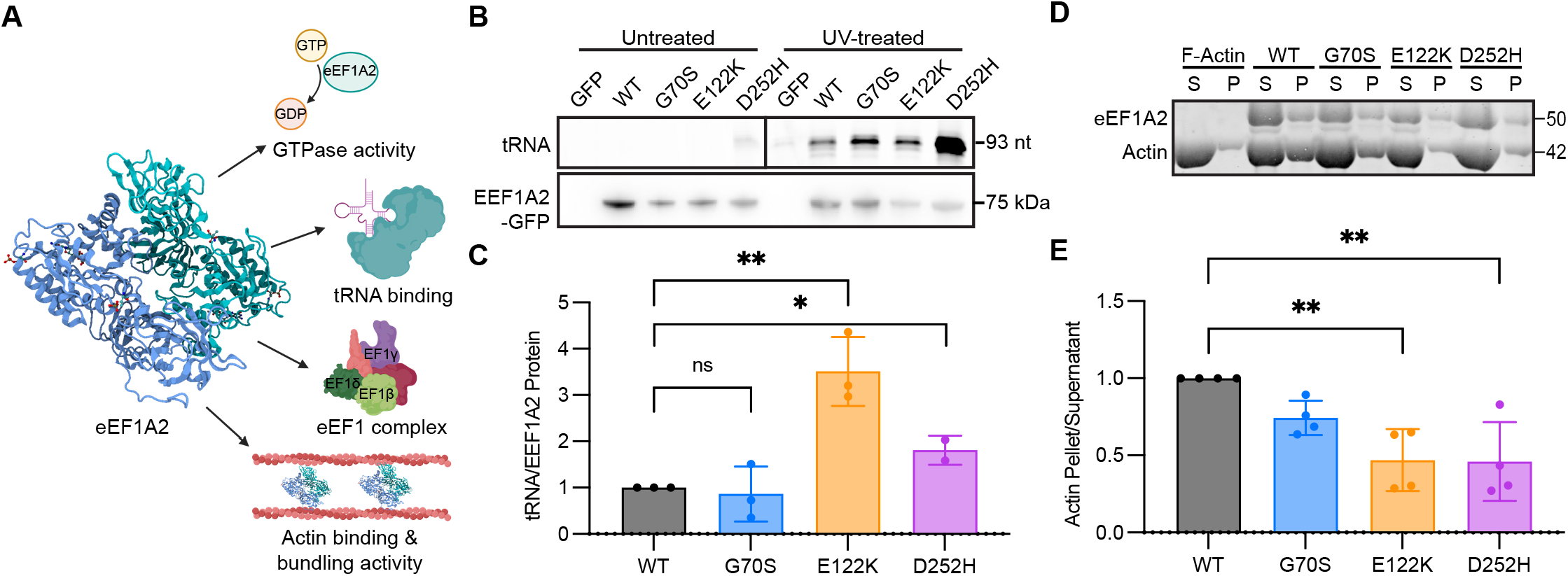
*EEF1A2* mutations alter tRNA binding and actin bundling activity A) Schematic of four eEF1A2 functions characterized in this study B) Representative denaturing RNA gel of total tRNA bound to eEF1A2-tRNA complexes immunoprecipitated from HEK293 cell lysates transfected with GFP-tagged *EEF1A2* or GFP-tagged G70S, E122K, and D252H mutants. Lower panel is a western blot of the immunoprecipitated complexes probing with an EEF1A2 antibody C) Quantification of total tRNA bound to eEF1A2-tRNA complexes immunoprecipitated from HEK293 cell lysates transfected with GFP-tagged eEF1A2 (WT)or GFP-tagged G70S, E122K, and D252H mutants. Data are normalized to wild-type EEF1A2 and represents the mean of three individual experiments ± standard deviation. One-way ANOVA with Tukey’s multiple comparison test *P<0.01, **P<0.009 D) F-actin bundling assay using flag-tag eEF1A2 proteins. Supernatant (S) and pellet (P) fractions were separated using a 4-12% bis-tris SDS-PAGE gel and stained with revert total protein stain (Licor). E)Quantification of F-actin bundling experiments with purified Flag-tag eEF1A2 or G70S, E122K, and D252H mutants. Data is the ratio of actin in the supernatant to the pellet and are normalized to wild-type eEF1A2 and represents the mean of four individual experiments ± standard deviation. One-way ANOVA with Tukey’s multiple comparison test. **P<0.005

Similarly, the *EEF1A2* mutations were found to decrease actin bundling activity. Cytoskeletal dynamics via actin binding and bundling is critical for the formation and stabilization of neuronal sub-compartments such as dendrites and spines (24). Given that eEF1A2 possesses an actin binding domain and can bundle actin filaments, we postulated that actin bundling activity might be altered by the disease-causing *EEF1A2* mutations. Purified Flag-tag eEF1A2 protein was incubated with filamentous actin and subjected to low-speed centrifugation. We found that both the E122K and D252H mutants significantly decreased actin bundling activity, whereas the G70S exhibited a small (∼15%) but nonsignificant decrease in bundling activity. This finding indicates that autism- and epilepsy-causing mutations in *EEF1A2* disrupt its actin bundling activity leading to alterations in the actin cytoskeleton in neurons.

## DISCUSSION

Here, we set out to characterize how mutations in *EEF1A2* result in functional changes to its protein product in an effort to uncover how these mutations lead to disease. The high penetrance and the severity of the neurological abnormalities in patients with *EEF1A2* mutations indicates its importance for neurological function and brain homeostasis. Consistent with these observations, we show that the three most common disease-causing *EEF1A2* mutations (G70S, E122K, and D252H) decrease de novo protein synthesis in both HEK293 cells and mouse cortical neurons. The mutation-induced decreases in protein synthesis appear to be driven by a decrease in the rate of peptide elongation. The GTPase activity of eEF1A2 was unaltered by these mutations, despite the proximity of some them to the active site. Notably, we found that all three disease-causing mutations increase the tRNA-binding capacity of eEF1A2. We hypothesize that the mutation-induced increase in tRNA binding results in a broad sequestration of all tRNAs and reduces tRNA availability, thereby reducing the rate of elongation. We also demonstrate that *EEF1A2* mutations decrease actin bundling activity disrupting the connection between the actin cytoskeleton and mRNA translation. We posit that these changes in the function of eEF1A2 lead to alterations in neuronal morphology, thereby changing neuronal connectivity and activity.

We have established that the autism- and epilepsy-associated mutations in *EEF1A2* are not simply loss of function mutations, but act via a gain of function by altering several aspects of eEF1A2 function, which then disrupts neuronal homeostasis. These findings are in line with previous studies that have examined the loss of one copy of *Eef1a2* and *EEF1A2* in both mouse and human, respectively. In *wasted* mice, loss of only one copy of *Eef1a2* does not result in either neurological or gait abnormalities, and the mice develop and age normally, exhibiting no aberrations in learning or memory (21). In humans, *EEF1A2* is among the genes deleted in benign familial neonatal epilepsy (BFNE), a disorder caused by a microdeletion of chromosome 20q13.3 (25). The epilepsy observed in this disorder has been attributed to the loss of *KCNQ2*, the gene adjacent to *EEF1A2*, which encodes the Kv7.2 subunit of potassium channels (26, 27). Interestingly, early in life these patients develop seizures that respond to epileptic drugs and eventually become seizure-free. Moreover, they lack any other neurological abnormalities and reach normal developmental milestones. In contrast, patients with *EEF1A2* mutations develop severe drug-resistant seizures and exhibit global developmental delay. Taken together, this evidence suggests haploinsufficiency of *EEF1A2* is not sufficient to cause either severe developmental or long-lasting neurological abnormalities.

Despite these previous observations, the link between *EEF1A2* mutations and the resulting neurological aberrations is not clear. Two possible hypotheses arise from our findings in this study. The first is that a loss of neuronal homeostasis is due to slowed or lack of production of neuronal proteins. Neurons have a large bias towards very long transcripts (28), so it is possible that the slowed elongation rate caused by the *EEF1A2* mutations results in changes in expression of neuronal specific genes and alterations to development, while sparing the translation of shorter transcripts. It has also been shown in other models of ASD that decreased translational efficiency on full length isoforms of risk genes underlies some of the pathogenesis of ASD (29). Further studies, using polysome and/or ribosome profiling should be able to rigorously test this hypothesis. An alternative hypothesis is based on the recent findings of Mendoza et al., (2021), which elegantly demonstrated the dissociation of eEF1A2 from its translation and actin-binding partners after phosphorylation in response to neuronal activity. The results of this study suggest that phosphorylation of eEF1A2 helps to coordinate translation with spine remodeling and overall synaptic plasticity. Although none of the *EEF1A2* mutations found in patients exist at the phosphorylation site of eEF1A2, the mutations could be mimicking phosphorylated eEF1A2, to reduce translation and actin bundling activity as we have observed here. In addition, mutations in *EEF1A2* may affect synaptic plasticity, a key neuronal process critical for learning and memory, which is known to be impaired in individuals with intellectual disability and many forms of ASD (30).

One limitation of our studies lies in the utilization of overexpression of eEF1A2 mutant proteins. Although eEF1A2 is highly expressed and gene dosage does not seem to affect cellular health, one cannot discount that artifacts may arise in our model system. In addition, *in vitro* assays assessing eEF1A2 functions are critical for understanding how the disease-causing mutations affect protein function, but are only approximations of what may be actually occurring *in situ*. Despite these limitations, we believe that our findings will open new avenues to understanding how autism- and epilepsy-associated mutations in *EEF1A2* result in disease and further expand our understanding of the regulation of neuronal protein synthesis.

## MATERIALS AND METHODS

### Cell Culture

HEK293 cells (ATCC) were cultured in Dulbecco’s Modified Eagles Medium (Gibco) supplemented with 10% Fetal Bovine Serum (FBS) (Gibco), 2 mM Glutamax, and 50 U/mL penicillin-streptomycin. Primary neuronal cultures were derived from mouse embryonic pups (e16.5-e18.5) (31). Neurons were plated on either glass coverslips (300,000 cells per 12 well) or 6 well plates (1.5 million cells per well) coated with Poly-D-Lysine and laminin. Cells were maintained in Neurobasal Plus (Gibco) supplement with B27 plus, 0.5 mM Glutamax, and 50 U/ml penicillin-streptomycin. Half medium changes were performed every 3-4 days with fresh growth medium. All cultures were maintained in a humidified incubator at 37°C and 5% CO2

### Cloning and Site-directed Mutagenesis

The *EEF1A2* CDS was cloned into pcDNA3.1-FLAG (gift from Saunders Lab) using NEBuilder® HiFi DNA Assembly Cloning Kit (NEB) according to the protocol of the manufacturer. The plasmid and coding sequence fragment was amplified using the following primers:

FLAG Vector For 5’- CGGGCAAGTAGCTCGAGTGCATTCTATAGTGTCACCTAAATGCTAGAGCTCGCT-3’,

FLAG Vector Rev 5’- TTCTCCTTGCCCATGGTTCGTCCCTTGTCGTCATCGTCTTTG

TAGT-3’

EEF1A2 CDS For 5’- AAGACGATGACGACAAGGGACGAACCATGGGCAAGG AG-3’

EEF1A2 CDS Rev 5’- TTTAGGTGACACTATAGAATGCACTCGAGCTACTTGCCC -3’

*EEF1A2* mutations in both the CMV-NGFP-EEF1A2-Neo vector and the pcDNA3.1-CMV-Flag-EEF1A2 vector were generated using the QuikChange II site directed mutagenesis kit (Agilent) according to the manufacturers protocol using the following primers.

G70S For 5’- GGAGCGTGAGCGCAGCATCACCATCGA-3’

G70S Rev 5’- TCGATGGTGATGCTGCGCTCACGCTCC-3’

E122K For 5’- CGGGCGTGGGCAAGTTCGAGGCG-3’

E122K Rev 5’- CGCCTCGAACTTGCCCACGCCCG-3’

D252H For 5’- GCCTGCCGCTGCAGCACGTGTACAAGATT-3’

D252H Rev 5’- AATCTTGTACACGTGCTGCAGCGGCAGGC-3’

### SDS-PAGE and Western Blotting

For western blotting, 20ug of protein sample was loaded into each lane of a 4-12% Bis-Tris polyacrylamide gel followed by a transfer to a polyvinylidene difluoride (PVDF) membrane. Total protein on the membrane was measured a No-Stain Protein Labeling Reagent (Invitrogen) and quantified using ImageJ. Membranes then were blocked using 5% milk in tris-buffered saline (TBS) for one hour and followed with incubation overnight with following primary antibodies in 5% bovine serum albumin (Puromycin 1:2000 (Millipore), GAPDH 1:1000 (Cell Signaling Technologies), eEF1A2 1:1000 (Abcam) Pan-Actin 1:1000 (Sigma), GAPDH 1:1000 (Cell Signaling Technologies), eEF1B2 1:1000 (Proteintech), eEF1D 1:1000 (Abnova), eEF1G 1:1000, (Santa Cruz Biotechnology)). Membranes then were washed 3 times for 5 minutes each with TBS 0.2% Tween and followed by incubation with horseradish peroxidase-conjugated secondary antibodies in 5% dry milk for one hour. Immunoreactive bands were detected using chemiluminescence (Cytiva) using the ProteinSimple imaging system (Bio-Techne). Band intensities were quantified using ImageJ.

### SUnSET Assay in HEK293 Cells

HEK293 cells were transfected with Lipofectamine 3000 (Invitrogen) according to the protocol of the manufacturer and incubated for 24 hours. Cells then were treated with 1ug/ml puromycin for 30 minutes and washed once with phosphate buffered saline. Cells were lysed in RIPA buffer (Thermo) supplemented with protease inhibitors. Lysates were incubated for 30 minutes on ice and then cleared by centrifugation for 15 minutes at 20,000g at 4oC. Protein concentrations were measured using a bicinchoninic acid (BCA) kit (Pierce, Thermo Scientific) using bovine serum albumin as a standard.

### SUNRISE Assay in HEK293 Cells

Cells were plated in 96 well plates coated with poly-d-lysine and transfected with Lipofectamine 3000 for 24 hours according to the protocol of the manufacturer. Cells initially were treated with harringtonine (2 ug/ml) or harringtonine and emetine (200 mM) for various durations of time, followed by treatment with puromycin at 10 ug/ml for 10 minutes. Cells then were washed with PBS and fixed for 10 minutes with 4% paraformaldehyde. Following permeabilization with 0.2% Triton-X cells were blocked with 5% normal goat serum for 1 hour and then incubated with an anti-puromycin antibody overnight. Cells were washed with PBS with 0.1% Tween-20 and then incubated with an anti-mouse secondary antibody conjugated to Alex Fluor 647 (Invitrogen). After washing, the cells were incubated with DAPI (10 ng/ml) (Invitrogen) in PBS for 10 min. Fluorescence measurements were made on a SpectraMax ID3 (Molecular Devices). Total puromycin fluorescence signal in each well was normalized to total GFP or DAPI fluorescence in the corresponding well.

### SUnSET Assay in Mouse Neurons

Primary mouse neurons plated on coverslips in 12 well plates were transfected with Lipofectamine 2000 at DIV8 with 1 ug of plasmid according to the protocol of the manufacturer. Neurons were incubated in transfection solution for 3 hours and then washed twice with PBS and a one-to-one mixture of conditioned media and fresh media was returned. At DIV14 cells were treated with 0.5 ug/ml of puromycin for 10 minutes and then fixed with 4% paraformaldehyde. Neurons were permeabilized with 0.2% Triton X and then blocked for 1 hour with 5% normal goat serum and incubated with the following primary antibodies in blocking buffer overnight (Guinea Pig MAP2 1:1000 (Synaptic Systems), Chicken GFP 1:1000 (Aves Bio), Rabbit EEF1A2 1:500 (ProteinTech) Mouse Puromycin 1:1000 (Millipore)). Neurons were washed 3 times with PBS with 0.1% Tween-20 and were incubated for 1 hour in the following secondary antibodies diluted in blocking buffer (anti-guinea pig AF 405 1:750 (Abcam), anti-chicken AF 488 (Abcam), anti-rabbit AF 568 (Invitrogen), anti-mouse AF 647 (Invitrogen). Coverslips were mounted on slides using Prolong-Gold (Invitrogen) and imaged using a confocal (Leica SP5) using a 40x oil immersion objective. Z-stack images were taken at 1 um steps for a total of 15 um. Images were analyzed using a custom ImageJ macro.

### Sholl Analysis of Mouse Neurons

Primary mouse neurons plated on coverslips in 12 well plates were transfected with Lipofectamine 2000 at DIV8 with 500 ng of pmCherryN1 (gift from Elias Spiliotis) and 500 ng of pCMV-NGFP-EEF1A2-Neo for a total of 1 ug according to the protocol of the manufacturer. Neurons were incubated in transfection solution for 3 hours and then washed twice with PBS and a one-to-one mixture of conditioned media and fresh media was returned. At DIV14 cells were treated with 4% paraformaldehyde. Neurons were permeabilized with 0.2% Triton X and then blocked for 1 hour with 5% normal goat serum and then incubated with the following primary antibodies in blocking buffer overnight (Guinea Pig MAP2 1:1000 (Synaptic Systems), Chicken GFP 1:1000 (Aves Bio), Rabbit EEF1A2 1:500 (ProteinTech), Rat mCherry 1:1000 (Invitrogen)). Neurons were washed 3 times with PBS with 0.1% Tween-20 and were incubated for 1 hour in the following secondary antibodies diluted in blocking buffer (Anti-Guinea Pig AF 405 1:750 (Abcam), Anti-Chicken AF 488 (Abcam), Anti-Rat AF 568 (Invitrogen), Anti-Rabbit AF 647 (Invitrogen)). Coverslips were mounted on slides using Prolong-Gold (Invitrogen) and imaged using a confocal (Leica SP5) using a 40x oil immersion objective. Z-stack images were taken at 1 um steps for a total of 15 um to completely capture the neuronal arbors. Images were analyzed using a custom ImageJ macro.

### FLAG Tag Purification

All FLAG-EEF1A2 variants were transfected into HEK293 cells and incubated for 48 hours. Cells were lysed in lysis buffer (50 mM Tris-HCl, pH 7.4, with 150 mM NaCl, 1 mM EDTA, and 1% Triton™ X-100) and cleared lysates were added to anti-FLAG magnetic beads (Sigma). Protein bound beads were washed with 3 washes with Tris-buffered saline (TBS) (50 mM Tris-HCl, 150 mM NaCl, pH 7.4) followed by 3 stringent washes with wash buffer (50 mM Tris-HCl, 500 mM NaCl, pH 7.4, 1% Tween-20). Beads were eluted using 3x Flag peptide (100ug/ml) in TBS. Purified protein then was concentrated with Pierce™ Protein Concentrators PES, 30K MWCO.

### Actin Bundling Assay

Actin bundling assays were carried out using the Actin Binding Protein Spin-Down Assay Biochem Kit (Cytoskeleton Inc.) according to protocol of the manufacturer.

### tRNA Binding Assay

Total tRNA binding was performed as described previously (32, 33). HEK293 cells transfected with either GFP, GFP-tagged eEF1A2, G70S, E122K, or D252H mutants were subjected to UV crosslinking with a BIO-RAD GS Gene linker set to 254 nm at 150 mJ/cm2 radiation. Cells then were lysed with 20 mM Tris-HCl, pH 7.4, 15 mM NaCl, 1% NP-40, 0.1% Triton® X-100, 1× HALT Protease Inhibitor Cocktail (Thermo). Cells that were not treated with UV light were lysed in parallel. eEF1A2-tRNA complexes were immunoprecipitated with GFP-TRAP magnetic beads (Proteintech). tRNAs were eluted from the beads using a hot-acid phenol extraction. tRNAs were ligated to a Cy3-labelled fluorescent stem-loop RNA/DNA oligonucleotide and subjected to gel electrophoresis using a 10% TBE-UREA denaturing-polyacrylamide gel (Invitrogen). tRNA was visualized using a Licor Odyssey imaging system.

### GTPase Activity Assay

Intrinsic GTPase activity was measured using a GTPase Assay Kit (Abcam) according to the protocol of the manufacturer. Absorbance was measured using a Spectramax ID3 plate reader (Molecular Devices)

### Co-Immunoprecipitation of Elongation Factor 1 Complex

HEK293 cells transfected with GFP or GFP-tagged EEF1A2, G70S, E122K, and D252H were lysed using co-IP lysis buffer (10 mM Tris/Cl pH 7.5, 150 mM NaCl, 0.5 mM EDTA, 2 mM MgCl2, 1 mM DTT, 0.5 % IGEPAL-630, 1x Halt Protease Inhibitor Cocktail). Cleared lysates (centrifugation at 20,000g for 15 minutes at 4C), were applied to GFP-TRAP magnetic beads (Proteintech), washed three times with wash buffer (10 mM Tris/Cl pH 7.5, 150 mM NaCl, 0.5 mM EDTA, 2 mM MgCl2, 1 mM DTT), and eluted with 1x SDS-polyacrylamide gel electrophoresis (PAGE) sample buffer and subjected to SDS-PAGE.

## Acknowledgments

This work was supported by NICHD F30 HD103360 (MSM), NINDS R35 NS122316 (EK), and US DOD W81XWH-21-1-0247 (EK). We would like to thank Sarah Reyman for help with optimization of actin bundling experiments and Emily Jang for help with figure preparation. We would also like to thank the members of the Klann laboratory for helpful insights and feedback. Schematics in Figure 1, 2 and 4 were created using biorender.com. Representative image panels in Figure 2 and 3 were created with QuickFigures (34).

## Author contributions

MSM and EK designed research. MSM preformed research and analyzed data. MSM and EK wrote the paper.

## Data and materials availability

All data are available in the main text or the supplementary materials.

