## Supplemental Figures for "Autism- and epilepsy-associated *EEF1A2* mutations lead to translational dysfunction and altered actin bundling"

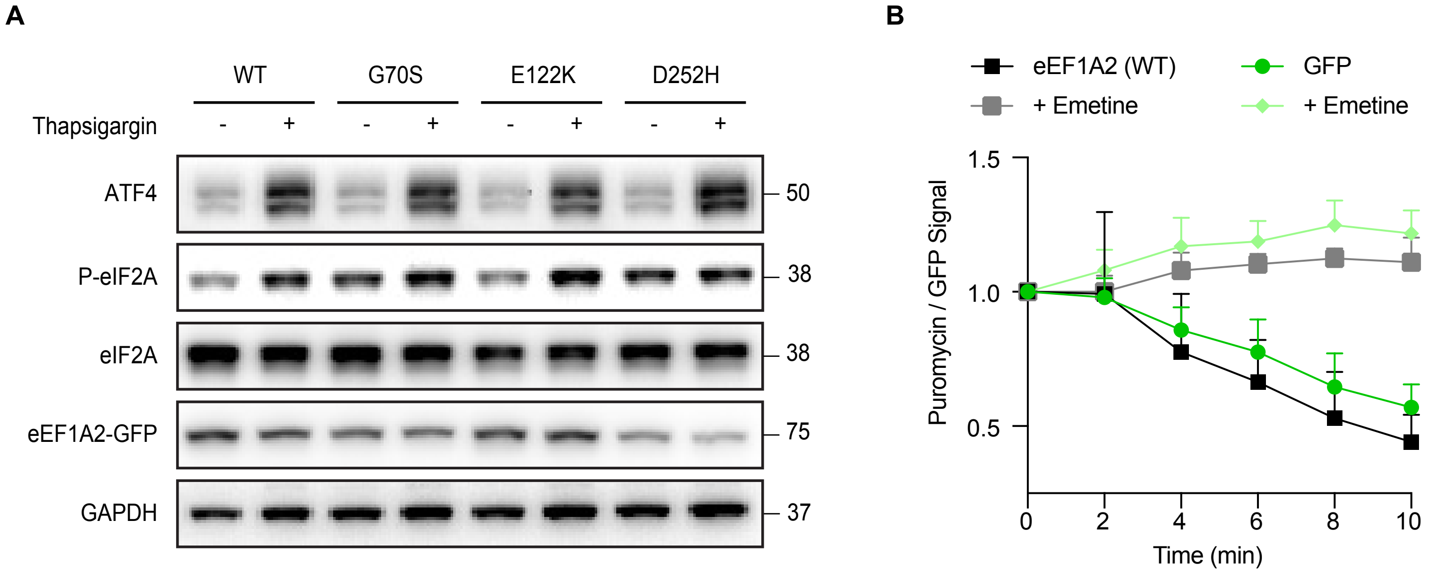


Supplemental Figure 1) *EEF1A2* mutations decrease de novo protein synthesis and elongation rates in HEK293 cells.

A) Representative western blot for markers of integrated stress response, ATF4 and phosphorylated-EIF2a. HEK293 cells transfected with wild-type EEF1A2, or G70S, E122K, D252H mutants.
B) SuNRISE in HEK293 cells transfected with wild-type EEF1A2 or GFP control. In addition to harringtonine treatment, emetine (200uM) was added at the same time as harringtonine.

### Supplementary Figure 1

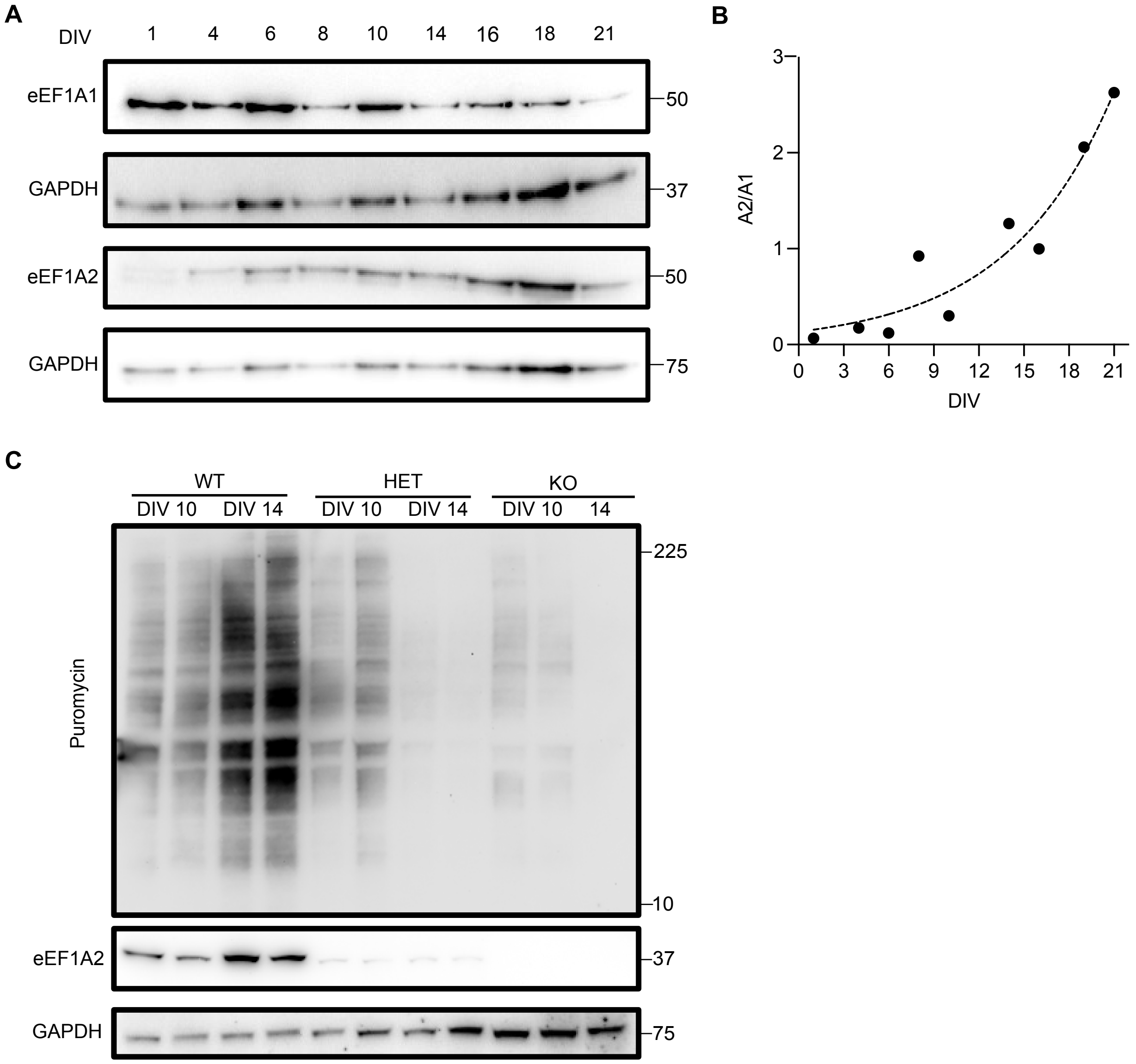


Supplemental Figure 2) eEF1 Isoform expression and proteins synthesis levels of *Eef1a2* null, *Eef1a2* heterozygous, and wild type cortical neurons in culture

A) Representative western blot for eEF1A1 and eEF1A2 expression over primary mouse neuron development
B) Quantification of the eEF1A2 to eEF1A1 expression ratio over the course primary mouse neuron development
C) SUnSET Assay in EEF1A2 +/+, EEF1A2 +/-, EEF1A2 -/- primary mouse neurons at DIV 10 and DIV 14.

### Supplementary Figure 2

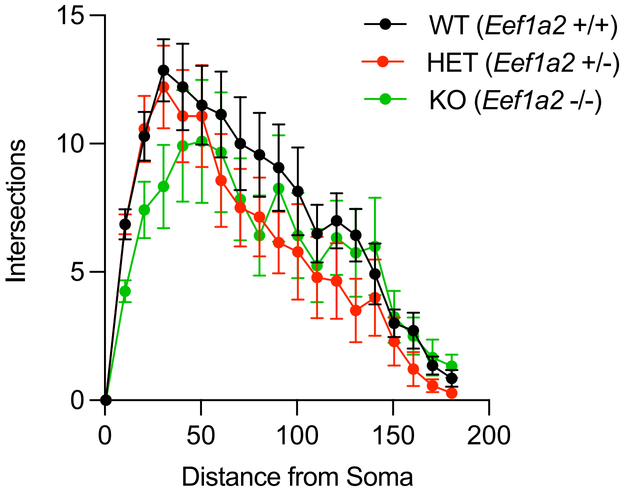


Supplement Figure 3) Sholl analysis in *Eef1a2* +/+, *Eef1a2* +/-, *Eef1a2* -/- primary mouse neurons transfected with mCherry at DIV 14.

#### Supplemental Figure 3

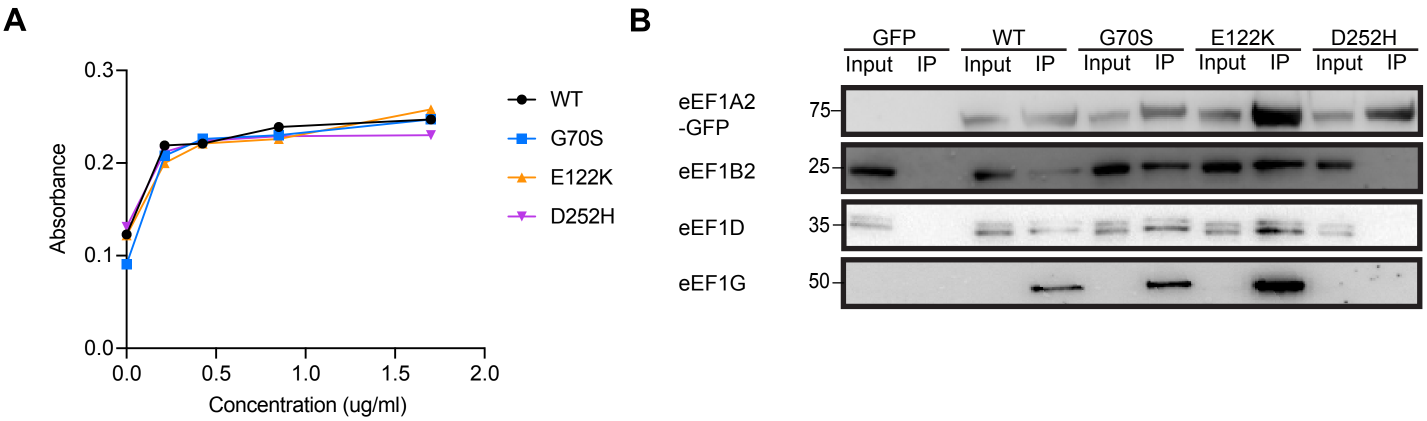


Supplemental Figure 4) In vitro characterization of eEF1A2 functions

A) Intrinsic GTPase activity of Flag-tag purified wild type EEF1A2 and G70S, E122K, D252H mutants using PiColorLock based GTPase activity assay.
B) Co-Immunoprecipitation GFP-tagged EEF1A2 complexes from HEK293 cells transfected with GFP or GFP-tagged EEF1A2, G70S, E122K, and D252H. Immunoblots of IP were probed with EEF1A2, EEF1B2, EEF1D and EEF1G. Experiment was repeated three times

### Supplemental Figure 4
